# Bio-DIA: A web-based tool for data and algorithms integration

**DOI:** 10.1101/2019.12.13.875666

**Authors:** Thiago Dantas, Vandeclécio Lira da Silva, André Faustino Fonseca, Diego A A Morais, Alberto Signoretti, Wilfredo Blanco

**Author notes:** Bioinformatics Multidisciplinary Environment (BioME), Federal University of Rio Grande do Norte (UFRN), Av. Odilon Gomes de Lima 1722, Natal/RN, CEP: 59078-400, Fone: 55 (84) 3342-2216. Computer Science Department, State University of Rio Grande do Norte (UERN – Natal), Av. Dr. João Medeiros Filho, 3419 - Natal/RN CEP: 59.120-200, Fone: 55 (84) 3207-8789/3207-2889.

## Abstract

Data science is historically a complex field, not only because of the huge amount of data and its variety of formats, but also because the necessity of collaboration between several specialists to retrieve valuable information. In this context, we created Bio-DIA, an online software to build data science workflow process focused in the integration of data and algorithms. Bio-DIA also facilitates the reusability of information/results obtained in previous process without the need of specific skills from the computer science field. The software was created with Angular at the front-end, Django at the back-end together with Spark to handle and process a variety of big data formats. The workflow/project is specified through XML file. Bio-DIA application facilitated the collaboration among users, allowing researcher’s groups to share data, scripts and information.

**Availability:** https://ucrania.imd.ufrn.br/biodia-app/. Login: bioguest, password: welcome123.

## Introduction

A data science process relies on a sequence of steps (pipeline) in which data is explored, integrated, analyzed, and processed (modeled, classified, etc), producing results that support (or not) to a given question/hypothesis. Several scientific fields are considered data-intensive, including biomedical sciences (Holzinger, Dehmer, and Jurisica 2014), an area where data science processes deals with a huge volume of data, coming from numerous sources and in distinct formats and levels of complexity. These issues have become a problem in biomedical research as well. Data from different formats and origins, such as symptoms, medical record information, clinical and laboratory data, medical images, genetic information, electrophysiological data, etc. tend to be poorly structured and mainly treated individually; as a result, studies usually are unable to integrate them and find possible relations.

To execute a given data science process in biomedical sciences, a group of professionals is usually composed by members from different areas (ex: biologists, mathematicians, computational scientists, statisticians, among others). They must collaborate to accomplish their goals and among other responsibilities, they must also decide or chose, within a vast domain of computer environments, languages and frameworks that are suitable for their necessities. This decision usually is taken based on individual knowledge or preference, that most of the time, it is not the best for the group or the tasks to be executed. Moreover, building upon previous computational experiment is almost mandatory in a collaborative environment. Research teams from all over the globe share source codes, data, and results in order to continue developing new projects. Therefore, reusability and reproducibility compose the essential building blocks in this type of scientific enterprise.

Under the described scenario, some issues are important to carry out a collaborative project, including: (1) data integration, (2) algorithms integration and (3) reusability.

This study proposes a web-based tool to facilitate the construction of data science process pipelines, called Bio-DIA, which combine big data exploration, integration and analyses. The results obtained by the built pipelines can be reused to replicate studies, facilitating the testing of new hypothesis and questions. This platform will facilitate the data, script and knowledge sharing among the scientific community.

We studied several software that work based on the three major points just mentioned. We found four software: Splunk, Galaxy, Kepler, and Knime. All of them have basically the same purpose, to built a data science process, focusing on data integration and analysis. Below a brief summary of these software.

Splunk (Stearley, Corwell, and Lord 2010) is an online platform that can connect with all type of data: sensors, structure data, NoSQL; through the use of Hadoop’s ecosystem. However, a subscription is required to use all features available in Splunk and the free version is very limited for big data analysis because they provide only 500MB for data storage.

Galaxy (Goecks, Nekrutenko, and Taylor 2010) is an open-source tool that was built for biomedical research. Any user can build new features to fill some personal need and it is also possible to use common functions like sort, select, join, or even specific functions, like the ones for Next Generation Sequencing (NGS). The major problem with Galaxy is the lack of documentation since any user can build new features, which makes the management of the system quite complex.

Kepler (Ludäscher et al. 2006) is an open-source Multi-platform desktop application, whose main goal is the creation of data science workflows to the research community in general. It is an open-source application, allowing the creation of your own modules (BioKepler (Altintas 2011) is an example). As KEPLER is an offline tool, the collaboration among researchers is more difficult.

Knime (Beisken et al. 2013) is also an open-source desktop tool (available off and online) with the objective to find, organize and mine data. The tool is built using Hadoop and Spark and contains more than 1500 modules and plenty of examples for starters. The desktop version is free to use, allowing the creation of personal features.

Bio-DIA was compared with these tools and some advantages were noticed: (1) totally free to use if compared with Splunk and Knime; (2) the use of xml to built data science process simplify the way to manipulate and share information/results if compared with the steep learning curve needed to used Galaxy; (3) since Bio-DIA is an online tool, only an internet browser is needed to executed data science process, without the necessity to install any additional software in the local machine, as Kepler does.

### Basic Structure of the Application

Bio-DIA is a web-based tool whose main goal is to provide an efficient platform for the integration of data and algorithms. Its front-end was developed using Angular 4, and the back-end uses Python programing language with Django framework as well as Spark. The use of Angular and Django allowed software scalability without loss of performance; and Spark permitted Bio-DIA to have a good scalability in the hardware side, being also capable to work with multiple types of data (Figure 1).

**Figure 1:**
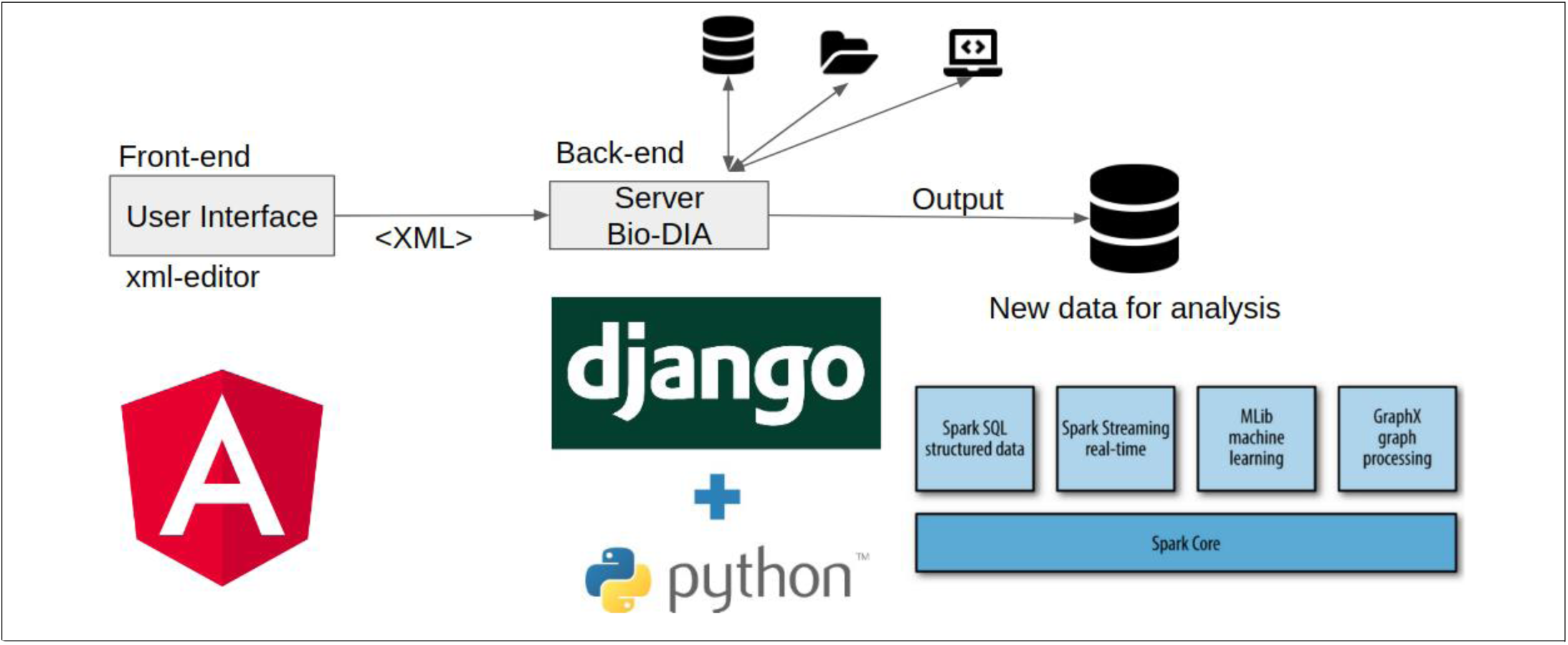
Bio-DIA architecture and technologies. The front-end was developed with Angular. A simple xml-editor was developed, allowing the user to create and submit his XML projects. For more details about the user interface, see **Supplementary Section 5**. After the XML is created, the front-end will send the xml file to the back-end (server) to be processed. In case the project requires any data, script or other project, Bio-DIA will carry out its execution. Once the project is finished, outputs are usually generated for further analysis. All the back-end was developed under Python/Django framework and Spark libraries as big data processing.

The BIO-DIA application is able to read and process (Selection, Projection, Filters, etc) several structured data types such as: text formatted files (tsv, cvs) and tables in databases (MySQL and PostgreSQL). The application is also able to load and run scripts (source codes) from different programming languages, such as Python, R, Perl and Shell.

Bio-DIA uses XML format to specify a data science workflow. The users are able to create a XML file (a “project” from now on) defining the workflow steps using several tags, such as: <data>, <select> and <save> to load, process and save data, respectively; the <algorithm> tag allows the loading and execution of scripts (see **Supplemental Section 1: Tags and attributes definition and description**). The data saved for a project can be used in another project using the tag <sub_project> (**Supplementary Section 6: Figure S9**).

Scripts usually need parameters to customize their execution. Using two embedded tags in the <algorithm> tags, Bio-DIA is able to send parameters to the script in two different ways. The first one is by using the <value> tag (**Supplementary Figure S8, line 7**), which is used to pass static values like numbers, strings or list of them (e.g.: a,b,c or 1,2,3); the second one is the <inter_data> tag (**Supplementary Figure S8, line 6**) used to reference data that was previously crated/generated in the current project. This is done through the value of the ***id*** typo, defined usually in the tags <data>, <select> or <sub_project>. This tag structure creates a standard scripting interface that allows a different code to be suitable for use by different researchers without them having to know computer coding.

A script executed inside a Bio-DIA project usually reads/writes (input/output) data or files. In order to control these events, Bio-DIA automatically add four (4) hidden parameters to decide where the data will be read or written. Therefore, the scripts have access to these parameters, which are named as: *DIR, ID, XMLID and PROJECT*; and they are sent in this order (more details, see **Supplementary Section 2**).

Bio-DIA was tested under a model HP Proliant DL386-G6 (AMD Opteron 243 with 12 cores) server, with 32 GB of RAM, and hard-disk space of 292GB.

## Results

In this section, we present 3’ projects in order to verify the results of Bio-DIA software according with the original pipeline executed. Although it could be use data from any science field, we focused on case studies applied to Bioinformatics field. First, we will present 2 projects that are usually very used on the Bioinformatics field, and finally we reproduce a previous work published by our group in (Silva et al 2015), that also used the two pipelines previously mentioned. We checked all the uploading, processing and writing events, comparing the results of the original code source.

### Case 01: Identifying conflicting genes names (Gene identifiers standardization)

This case was built to address the inconsistency regarding the gene identification (*id*) value in different databases, such as id conversions between Entrez (Maglott et al. 2005), Ensembl (Flicek et al. 2013), HUGO Symbol (Eyre et al. 2006) and RefSeq (Pruitt, Tatusova, and Maglott 2005). The project was created to confer and identify values discrepancy, which can be a time-consuming activity, delaying significantly the projects during data integration and cross-database analysis. This bottleneck is often related to a large data volume and data version, producing false associations and possible misinterpretations. Nowadays, several tools are available to solve this problem, like MyGene (Xin et al. 2015) and Biomart (Durinck et al. 2005). In this context, we illustrate a protocol to bypass this problem using Bio-DIA XML syntax (**Figure 2**).

**Figure 2:**
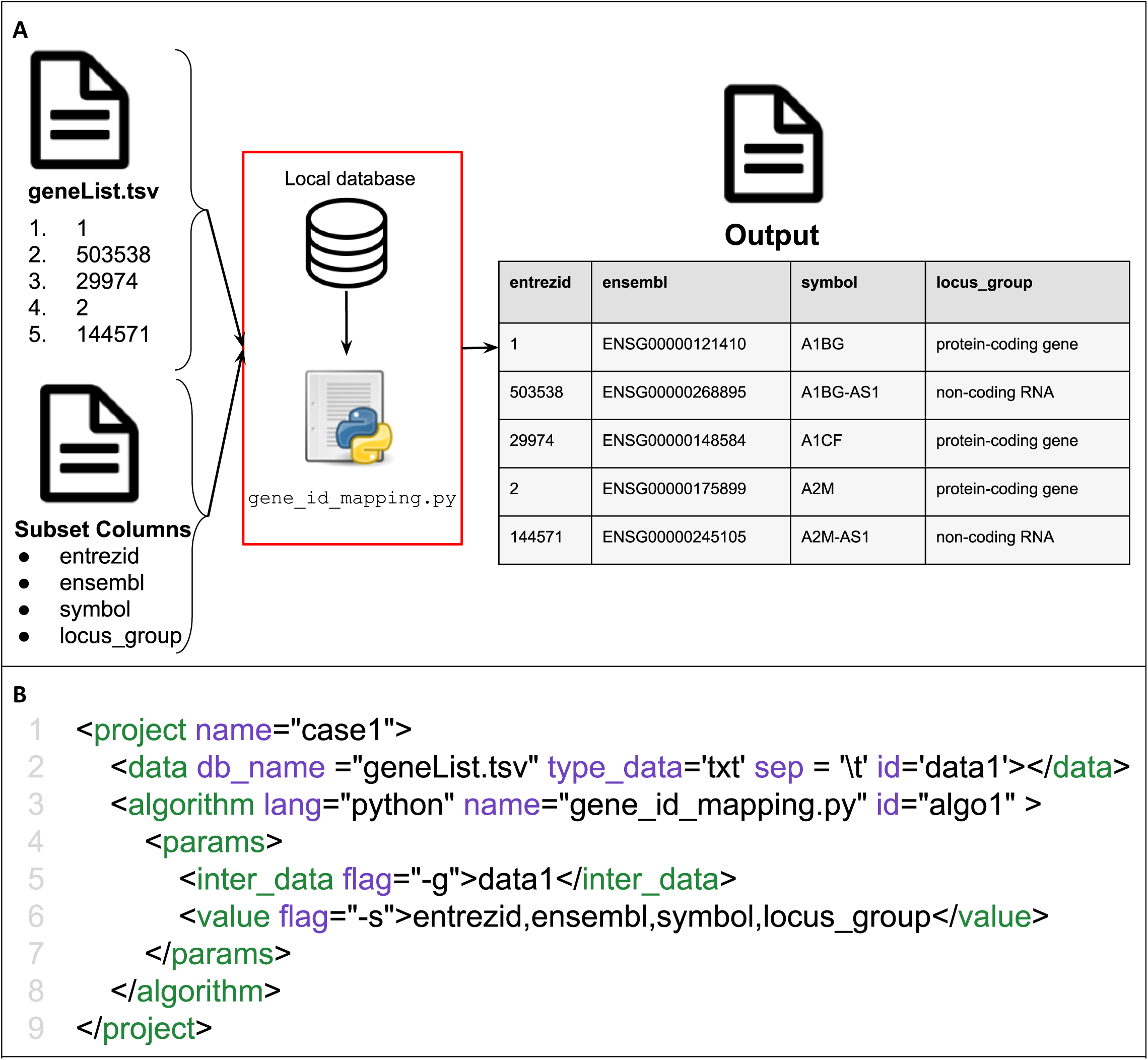
Gene identifiers standardization protocol. (A) The workflow schema of the pipeline is shown, in which a python script algorithm takes two external inputs: a list of gene identifiers (geneList.tsv), and the subset of columns to produces a TSV file (Output). A local database file is consulted by the script to show the distinct ID types (such as: HUGO Symbol, Ensembl Gene, Transcript RefSeq, and NCBI Entrez) from the selected inputs (geneList.tsv). The red box delimited shows an internal input that the script will be able to access. (B) XML script to execute the pipeline shown in A. Line 2, using <data> tag, is in charge to load the gene list input (geneList.tsv file seem in A). Line 3 defines a python-type scrip named “gene_id_mapping.py”. From line 4 to 7 are defined the parameters for the algorithm. The first parameter (line 5), is the gene list that is referenced by the ‘data1’ which was previously defined in line 2 as *id*. The second parameter (line 6), passed as values, is the subset columns.

### Case 02: Genes related with a tumor and platform

In the present case, the goal of the task is to search and retrieve a specific list of genes from TCGA (https://tcga-data.nci.nih.gov), an external database that stores data from roughly eleven thousand patients comprising 32 tumor types. The R script gets 3 parameters: the list of desired genes, the tumor name (in this case GBM), and the type of the filter (platform) that is ‘mrna’ in the present case. The output is a text file containing 5 columns (sample, gbm_tcga_rna_seq_v2_mrna, gbm_tcga_mrna, gbm_tcga_mrna_U133 and gene name), and their names will change according to the parameters passed. A workflow and the corresponding XML file of this case are shown in Figure 3.

**Figure 3:**
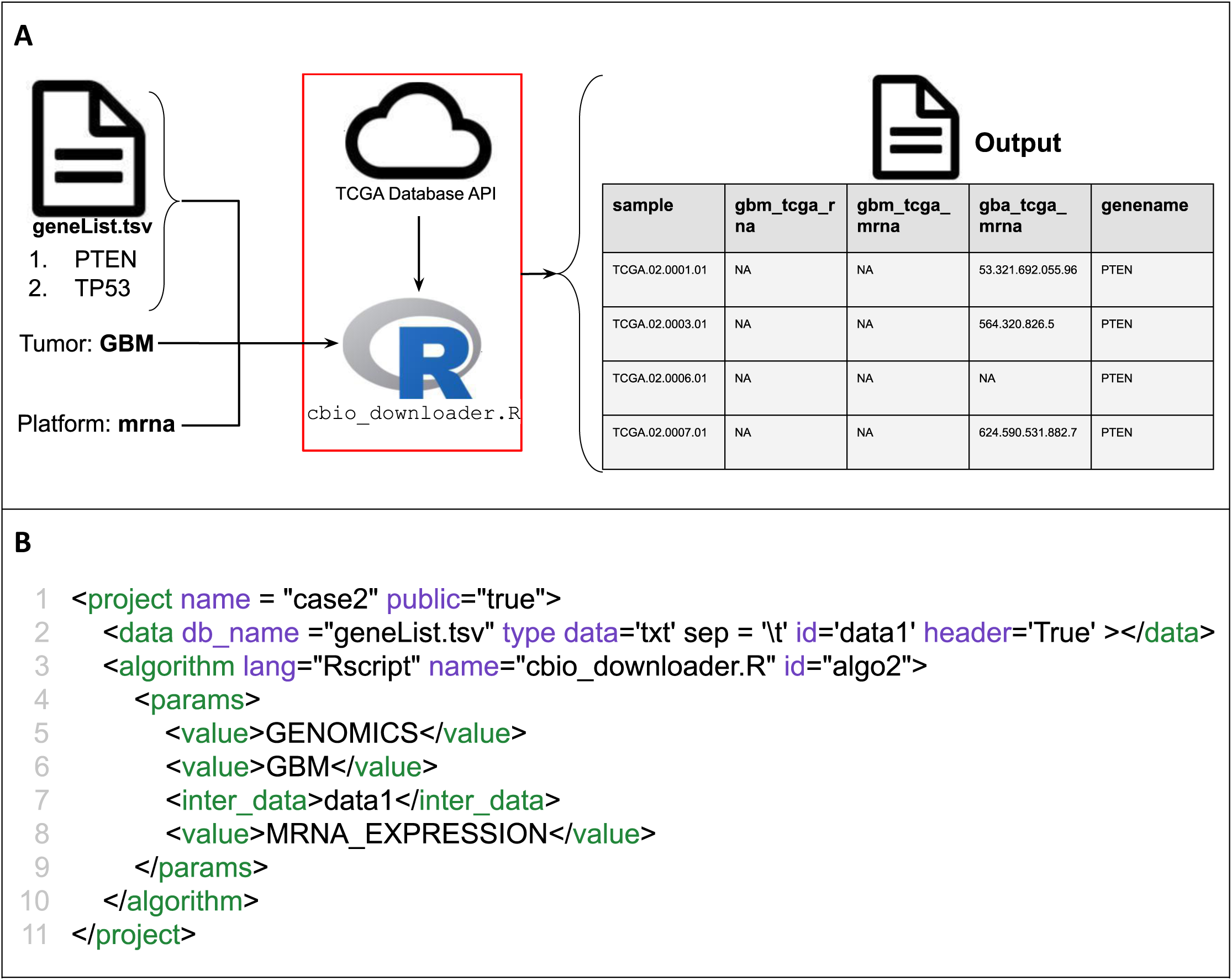
Pipeline to retrieve list of genes related with a tumor and platform. (A) The workflow schema with three inputs parameters: gene list, tumor name and platform. The R script (cbio_downloader.R) connects with the TCGA external database and retrieve a filtered list with the genes related with the tumor GBM(Glioblastoma) and mrna as platform. The red box is just delimiting the R algorithm context, which is in charge to connect and access TCGA. (B) XML script to execute the pipeline shown in A. The line 1 defines the project tag and its attributes (name and public, see **Supplementary Section 1**). The line 2 uses a <*data*> tag to load the gene list file (*db_name = “geneList.tsv”).* From lines 3-10 the R script is executed with three parameters (from lines 4 to 7): lines 5,6 and 8, through <value> tag, the ‘GENOMICS’, ‘GBM’ and ‘MRNA_EXPRESSION’ are passed as string values respectively; in line 7, through <inter_data> tag, the *data1* is passed, which is pointing to the list of genes referenced by the *id* created in line 2.

### Case 3: Cancer/testis (CT) project

The CT study was published by our group in (da Silva et al. 2017), in which the integration of gene expression and clinical data guided us to detect some CT genes that are associated to prognosis in different types of cancer. This study executed a genome-wide screen for CT genes using data from several databases; hence, we reproduced the original pipeline (Figure 4A) using the Bio-DIA XML script (Figure 4B, C) in order to replicate the results.

**Figure 4:**
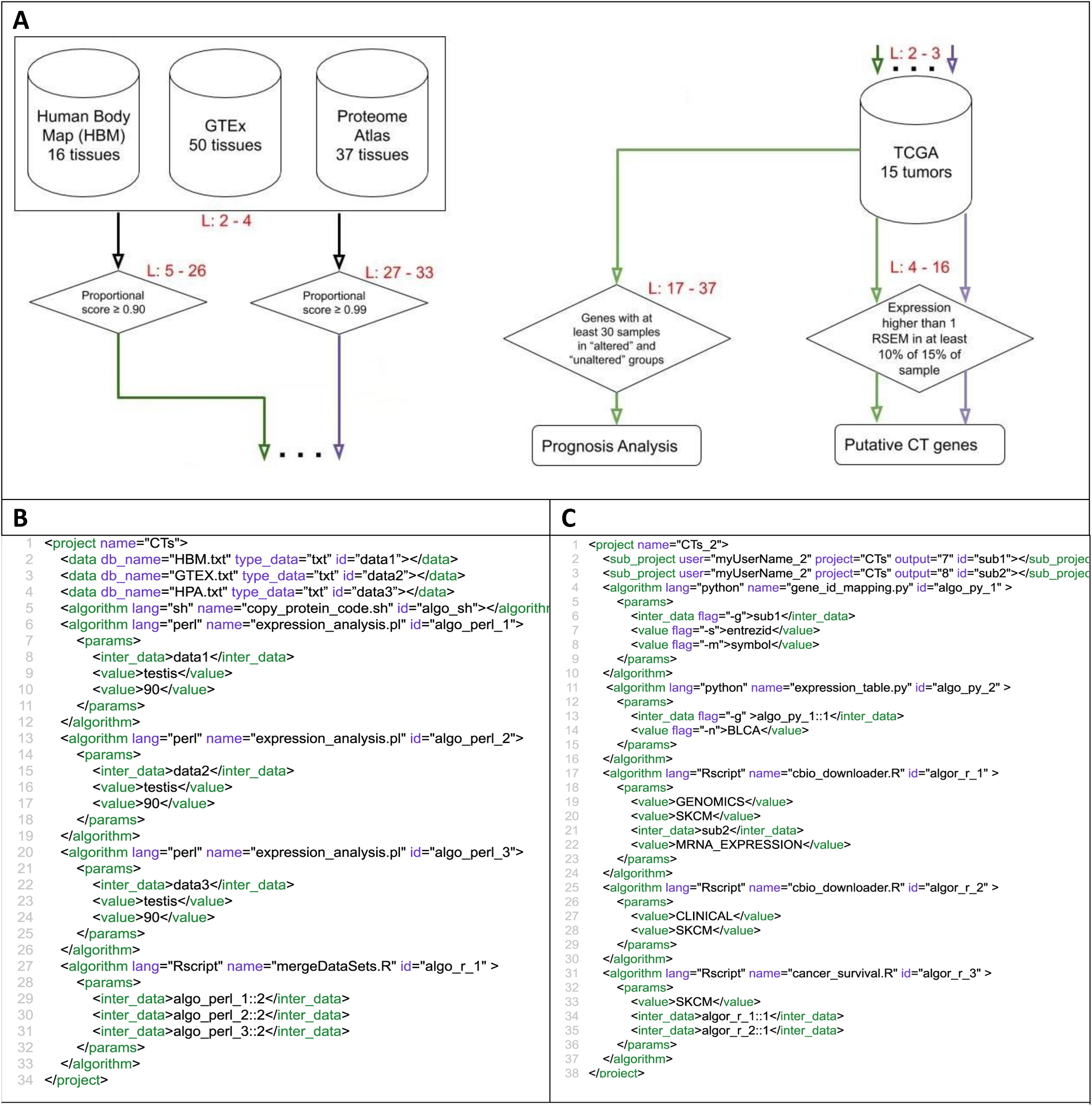
CT’s project built on Bio-DIA XML script. (A) Schematic pipeline of the original CT’s project, but divided in left and right side, to better explain its implementation in Bio-DIA projects. Red labels correspond to the XML lines in charge to execute each part (modified from Da Silva et al, 2017). (B) XML script lines that reproduce the first project (A left side) so-called ***CTs***. It starts accessing 3 databases using <data> tags (lines 2, 3 and 4 for HBM, GTEX and HPA databases respectively). In line 5, a script shell, named “copy_protein_code.sh”, is called using <algorithm> tag. This script is in charge to copy to the Bio-DIA folder (defined by DIR parameter) a list of genes from the human genome that are validated by the NCBI, making possible to get only protein-coding genes from these 3 databases. Subsequently (from line 6-26), a Perl source code, named “expression_anaysis.pl”, is called 3 times to filter the databases. Each call has three parameters: the database (using <inter_data> tag), the tissue and the cutoff. The last two arguments, both are passed by values (using <value> tag). The last step, an R source code is called with 3 parameters, which are the second output of each perl algorithm previous executed (using <inter_data> algo_perl_1::2</inter_data>). (C) XML script lines that reproduce the second project so-called ***CTs_2***. Through the <sub_project> tag (Line 2) and its attribute *output*, BIO-DIA is able to load the specific output 7 (listGeneName.txt) from the project already executed, ***CTs.*** *Then*, a python script (case1) is executed, receiving the <sub_project id=‘sub1’> and the parameters -s for subset columns and the ‘-m’ for the mapping type. After that, a new script is executed ‘expression_table.py’, receiving as parameter the first output generated(geneList) by the previous script ‘gene_id_expression.py’, and the second parameter ‘-n’ passing the tumor name (BLCA). The next script on line 17 – 30 is used the case 2, getting Genomics and Clinical data from the tumor (SKCM), finally, the algorithm in line 31, is receiving as parameters the data collected by the scripts(algo_r_1 and algo_r_2) with these outputs, it will be possible to make plots to check what genes are helping the patients to live longer.

The methodology applied on CTs study (da Silva et al. 2017) focuses on data integration from distinct sources. Three RNA-Seq databases from normal tissues were used to identify genes with expression bias to testis: the Human Body Map (HBM) (GEO accession: GSE30611), Genotype-Tissue Expression (GTEx) [DOI:10.1093/nar/gkv1045] and Human Proteome Atlas (HPA) [DOI:10.1126/science.1260419]. Normalized transcript level in each tissue was converted to a proportional score (transcript level in a tissue divided by the sum of levels in all tissues), and a threshold of at least 0.9 was used to select genes preferentially or exclusively expressed in testis (additionally, a more restrict threshold of 0.99 was performed).

In the next step from the pipeline, RNA-Seq data from TCGA with 6,221 tumor samples and 15 tumor types were used to identify genes significantly expressed in a given tumor, considering them as putative CT gene if it had a level of expression (cutoff threshold of RSEM >1) in at least 10% or 15% of all informative samples for a given tumor. Finally, the integration of gene expression, abundance of CD8+ cells infiltration and clinical data led us to identify dozens of CT genes associated with either good or poor prognosis observing the survival curve.

## Conclusions

We implemented an application to help the scientific community to overcome issues related with data/algorithm integration and replicability; with the main goal to provide a web-based tool to improve scientific collaboration. The Bio-DIA project/pipelines, specified by XML, intent to make easier the access to big data with different types of formats, execute algorithms (scripts) developed in different languages (R, python, Shell, and Perl) and reuse/share these results among projects.

Bio-DIA can be accessed via public site at https://ucrania.imd.ufrn.br/biodia-app/. We developed a simple graphic user interface (GUI), providing the necessary functionalities to manage (create, retrieve, edit, remove and execute) the projects and their results. More details and explanation of the GUI can be found in **Supplementary Section 5**. In further versions, a new GUI will be available to manage and automatically generate projects by only dragging-and-dropping graphic shapes, icons and their connections. Thus, knowledge of XML will no longer be required to use Bio-DIA, making its use even simpler for a heterogeneous data science team.

## References

Holzinger, Andreas, Matthias Dehmer, and Igor Jurisica. 2014. “Knowledge discovery and interactive data mining in bioinformatics-state-of-the-art, future challenges and research directions.” BMC bioinformatics 15 (6):I1.

Altintas, Ilkay. 2011. Distributed workflow-driven analysis of large-scale biological data using biokepler. Proceedings of the 2nd international workshop on Petascal data analytics: challenges and opportunities.

Beisken, Stephan, Thorsten Meinl, Bernd Wiswedel, Luis F de Figueiredo, Michael Berthold, and Christoph Steinbeck. 2013. KNIME-CDK: Workflow-driven cheminformatics. BMC bioinformatics 14 (1):257.

da Silva, Vandeclecio Lira, André Faustino Fonseca, Marbella Fonseca, Thayna Emilia da Silva, Ana Carolina Coelho, José Eduardo Kroll, Jorge Estefano Santana de Souza, Beatriz Stransky, Gustavo Antonio de Souza, and Sandro José de Souza. 2017. Genome-wide identification of cancer/testis genes and their association with prognosis in a pan-cancer analysis. Oncotarget 8 (54):92966.

Durinck, Steffen, Yves Moreau, Arek Kasprzyk, Sean Davis, Bart De Moor, Alvis Brazma, and Wolfgang Huber. 2005. BioMart and Bioconductor: a powerful link between biological databases and microarray data analysis. Bioinformatics 21 (16):3439–3440.

Eyre, Tina A., Fabrice Ducluzeau, Tam P. Sneddon, Sue Povey, Elspeth A. Bruford, and Michael J. Lush. 2006. The HUGO Gene Nomenclature Database, 2006 updates. Nucleic Acids Research 34 (suppl_1):D319-D321. doi: 10.1093/nar/gkj147

Flicek, Paul, M. Ridwan Amode, Daniel Barrell, Kathryn Beal, Konstantinos Billis, Simon Brent, Denise Carvalho-Silva, Peter Clapham, Guy Coates, Stephen Fitzgerald, Laurent Gil, Carlos García Girón, Leo Gordon, Thibaut Hourlier, Sarah Hunt, Nathan Johnson, Thomas Juettemann, Andreas K. Kähäri, Stephen Keenan, Eugene Kulesha, Fergal J. Martin, Thomas Maurel, William M. McLaren, Daniel N. Murphy, Rishi Nag, Bert Overduin, Miguel Pignatelli, Bethan Pritchard, Emily Pritchard, Harpreet S. Riat, Magali Ruffier, Daniel Sheppard, Kieron Taylor, Anja Thormann, Stephen J. Trevanion, Alessandro Vullo, Steven P. Wilder, Mark Wilson, Amonida Zadissa, Bronwen L. Aken, Ewan Birney, Fiona Cunningham, Jennifer Harrow, Javier Herrero, Tim J.P. Hubbard, Rhoda Kinsella, Matthieu Muffato, Anne Parker, Giulietta Spudich, Andy Yates, Daniel R. Zerbino, and Stephen M.J. Searle. 2013. Ensembl 2014. Nucleic Acids Research 42 (D1):D749-D755. doi: 10.1093/nar/gkt1196

Goecks, Jeremy, Anton Nekrutenko, and James Taylor. 2010. Galaxy: a comprehensive approach for supporting accessible, reproducible, and transparent computational research in the life sciences. Genome biology 11 (8):R86.

Ludäscher, Bertram, Ilkay Altintas, Chad Berkley, Dan Higgins, Efrat Jaeger, Matthew Jones, Edward A Lee, Jing Tao, and Yang Zhao. 2006. Scientific workflow management and the Kepler system. Concurrency and Computation: Practice and Experience 18 (10):1039–1065.

Maglott, Donna, Jim Ostell, Kim D. Pruitt, and Tatiana Tatusova. 2005. Entrez Gene: gene-centered information at NCBI. Nucleic Acids Research 33 (suppl_1):D54-D58. doi: 10.1093/nar/gki031

Pruitt, Kim D., Tatiana Tatusova, and Donna R. Maglott. 2005. NCBI Reference Sequence (RefSeq): a curated non-redundant sequence database of genomes, transcripts and proteins. Nucleic Acids Research 33 (suppl_1):D501-D504. doi: 10.1093/nar/gki025

Stearley, Jon, Sophia Corwell, and Ken Lord. 2010. Bridging the Gaps: Joining Information Sources with Splunk. SLAML.

Xin, Jiwen, Adam Mark, Cyrus Afrasiabi, Ginger Tsueng, Moritz Juchler, Nikhil Gopal, Gregory S. Stupp, Timothy E. Putman, Benjamin J. Ainscough, Obi L. Griffith, Ali Torkamani, Patricia L. Whetzel, Christopher J. Mungall, Sean D. Mooney, Andrew I. Su, and Chunlei Wu. 2015. MyGene.info and MyVariant.info: Gene and Variant Annotation Query Services. bioRxiv:035667. doi: 10.1101/035667

