## Supplementary Materials for "Bio-DIA: A web-based tool for data and algorithms integration"

+ 55 (84) 98729-0888

### Supplementary Section

#### Supplementary Section 1: XML tags and attributes

| Tag | Attributes | | Text | Child | Definition |
|---|---|---|---|---|---|
| | Required | Not Required | | | |
| <project> | name | None | None | All tag's below | Used to create a project. |
| <data> | id<br>db_table<br>type_data | db_name<br>sep<br>comment<br>header | None | No | Used to access some data format such as: csv, txt, tsv, and tables from MySQL and Postgre databases. |
| <sub_project> | id<br>user<br>project<br>output | None | None | No | Used to load output data from another project and used it in the current project. |
| <algorithm> | id<br>lang<br>name | None | None | Yes: <params> | Used to call some algorithm developed on specific script language, such as: Python, R, etc. |
| <params> | None | None | None | Yes: <inter_data> or <value> | Inform the arguments needed to execute the script. |
| <inter_data> | None | None | True | No | Used to link with some databases that exists inside the project. |
| <value> | None | None | True | No | Inform some value that the script need to execute. |
| <select> | id<br>from<br>get | filter<br>limit | None | No | Spark function to make a select on the database. |
| <save> | Store<br>Folder | None | None | No | Save something that will be executed on the project on your project folder. |

| Attribute | Definition | Examples |
|---|---|---|
| Id | Used to identify and reference a tag inside the XML. Its value is a string/text type format and is unique in the all XML project. | <data id=“myDATA”/></data><br><br><select from=“myDATA” /></select> |
| db_table | Used only inside the <data> tag to connect with ether a table from an database or text file. | <data db_table=”myFile.txt”></data> |
| db_name | Used to identify the database name to be connected. In order to open a table, the <db_table> attribute has to be specified. | <data db_name = “myDB” db_table=”myTable”> |
| type_data | It is used to inform which data type will be used, the types are: txt, sql and pg. The txt value is used to say that it will work with text files (txt, csv and tsv). The sql value is to say that it will work with MySQL(Community Edition ), and pg with Postgre. | <data db_name= “myFile.txt” type_data=”txt”></data><br><br><data db_name=“myDB” type_data=”sql”></data> |
| sep | To enter the file separator type, either comma separated values or \t(tab). | <data db_name= “myFile.txt” sep=”\t”></data><br><br><data db_name= “myFile.csv” sep=”,”></data> |
| comment | If it has comment lines to ignore, it must be specified the character (#,//,;). | <data db_name= “myFile.txt” comment=“#”></data><br><br><data db_name= “myFile.txt” comment=“;”></data> |
| header | If it is a text file, it must be informed if it has header passing the value True, if it exists or False. | <data db_name=“myFile.txt” header=“True”></data><br><br><data db_name= “myFile.txt” header=“False”></data> |
| user | Used to enter the name of the user providing the project outputs. | <sub_project user= "userName_id" ></sub_project> |
| project | Name of the project that the user has set to public for all users. | <sub_project user="userName_id" project="projectName"> </sub_project><br><sub_project user="userName_id" project="projectName"></sub_project> |
| output | Output number that tells which output the user wants to get for his own project. | <sub_project user="userName_id" project="projectName" output="output_number" ></sub_project><br><br><sub_project user="userName_id" project="projectName" output="outputNumber"></sub_project> |

| | | |
|---|---|---|
| from | Receive which will be the database will be used, the value of this attribute must be some ID of some tag of type <date> or <sub_project> | <data db_name="myDB" type_data= “sql” id=“data_1” ></data><br><br><select from=“data_1” ><select/> |
| get | It must be informed which columns should be returned in the query. | <select from=“data_1” get=“column_a, column_b”><select/> |
| filter | The filter condition must be passed to the query. | <select id=“select1” from=“data_1” get=“column_a, column_b” filter=“column_a like ‘test’ “><select/> |
| limit | Number of rows to be returned, if the attribute is missing, all rows will be returned. | <select id=“select1” from=“data_1” get=“column_a, column_b” filter=“column_a like ‘test’ “ limit=“50” ><select/> |
| inplace | If the value is True, the original data of the from attribute will be overwritten by the select query result, if False, will not change the original data, a new file will be created, to save this file, just need to pass the id of the <select> on the tag <save>. | <select id=“select1” from=“data_1” get=“column_a, column_b” filter=“column_a like ‘test’ “ limit=“50” inplace=“True” ><select/><br><br><select id=“select1” from=“data_1” get=“column_a, column_b” filter=“column_a like ‘test’ “ limit=“50” inplace=“False” ><select/> |
| Lang | Used to specify the programming language to be used (Rscript, python or perl). | <algorithm lang=“perl" ></algorithm><br><algorithm lang="Rscript" ></algorithm> |
| name | Name of the script. | <algorithm lang="perl" name="myCode.pl" ><br></algorithm> |
| Flag | This attribute can only be used in the <inter_data> or <value> tags, its functionality is to pass some argument to the script. | <params><br><inter_data flag=“-x”>select1</inter_data><br><value flag=“-y”>23,22</value><br></params> |

| | | |
|---|---|---|
| store | Name of the id tag inside the project that you want to save on your project folder. | <select id="select1" from="data_1" get="column_a, column_b" filter="column_a like 'test' " limit="50" inplace="False" ><select/><br><br><save store="select1" ></save> |
| folder | Name of the folder that the output will be saved, by default the folder name is the same of the project | <project name="projectName"><br><select id="select1" from="data_1" get="column_a, column_b" filter="column_a like 'test' " limit="50" inplace="False" ><select/><br><br><save store= "select1" folder="projectName"><br></save> </project> |

Scripts usually need parameters to customize their execution and the <parameter> tag is used to define them. Bio-DIA has two ways to send parameters to the script; the first one is using the <value> tag (Supplementary Figure S1, line 8), which is used to pass static values like numbers, strings, or list of them (e.g.: a,b,c or 1,2,3); the second one is the <inter_data> tag (Supplementary Figure S1, line 7) used to reference data that was previously crated/generated in the current project. This is done through the value of the *id* attribute, defined usually in the tags <data>, <select> or <sub_project>.

#### Supplementary Section 2: How Bio-DIA execute scripts

In Figure S1 depicts that Bio-DIA can execute scripts from languages (Shell, Python, Rscript or Perl) using the <algorithm> tag. In order to Bio-DIA be able to execute the scripts on the server, it will send add four (4) hidden parameters (DIR, ID, XMLID, PROJECT) mainly to control the reading and writing events (Figure S1). Consequently, the scripts have access to these parameters. Case the script does not need using flags as parameters, these hidden arguments will be added in the end, if the script need flags to send parameters, the developer must reserve the flag '-c' as default to receive these 4 hidden parameters.

The DIR parameter defines the directory path where Bio-DIA will load or save files; in other words, any access to the server hard disk memory will be done in this directory. All files that will be passed as <inter_data> the script may read it using the DIR. The ID parameter stores the user identification value, this ID it will be used to create a file to write everything that happen with the script (log file). A log file it will be a way to know what occurred inside the script. The log file is always named like ID_log.tsv and have 2 columns named as STATUS and MSG. The first column is to specify which is happening inside the script, like: [START], [RUN], [OUTPUTS], [ERROR] and [CONNECT], the second column is the message related with the status, like: "[START] The script just started" or "[CONNECT]  Connection made with success with the database".

The third parameter XMLID (script id on XML pipeline) is used if the script will write any output the procedure is: the output must be described on the log file, with the status [OUTPUTS]; the message in this status must be using this template XMLID::PROJECT_myOutput.csv.

The fourth parameter PROJECT, is to be concatenated with every output name created by the script, allowing Bio-DIA to recognize and control the outputs from the projects. The name pattern looks like PROJECT_myOutput.csv.

All these parameters along with the log file, allows Bio-DIA to know the outputs generated for the projects, consequentially other users re-use these outputs on their projects. Check the figure below to see an example and values of these four parameters.

**A**

| Parameter | Value |
|---|---|
| DIR | /home/biodia/project/ |
| ID | 1 |
| XMLID | algorithm1 |
| PROJECT | projectName |

**B**

Log file name: ID_log.tsv (1_log.tsv)

| STATUS | MSG |
|---|---|
| [START] | Start Script {ID:algorithm1}. |
| [RUN] | Trying to read geneList as input. |
| [RUN] | Connecting with database. |
| [OUTPUTS] | algorithm1::projectName_myOutput.csv |
| [END] | Finished. |

File save path: /home/biodia/project/
Outputs written:
1 log.tsv, projectName_myOutput.csv

**Figure S1:** Log file and hidden parameters example. (A) table with the four hidden parameters and values, that all scripts will receive before being executed, these parameters are an example of how the script in the Figure S1, line 4 will receive. (B) shows how the log file is build, and how the parameters are used and where the parameters must be placed following the procedures talked on Section (How to execute scripts on Bio-DIA).

#### Supplementary Section 3: Passing outputs from an algorithm to another one within the same project.

It is possible to pass an output from an algorithm to another one (as a parameter) in the same XML project. The Figure S2 shows an example of how to do this.

```
1  <project name="myPipeline">
2      <algorithm lang="python" name="mathSum.py" id="algo_py_1" >
3          <params>
4              <value flag="-x">1,2,3,4,5,6,7,8,9,10</value>
5          </params>
6      </algorithm>
7       <algorithm lang="python" name="mathPow.py" id="algo_py_2" >
8          <params>
9              <inter_data>algo_py_1::1</inter_data>
10             <value>3</value>
11         </params>
12     </algorithm>
13 </project>
```

**Figure S2.** Example how to access output among algorithms. The algorithm 'algo_py_1' is receiving a list of numbers as parameters to be added. The output will be saved in a file with the summation(Sum) of this list, the second algorithm 'algo_py_2' need two parameters, the first is the output created by the first algorithm 'algo_py_1', to access this output the syntax is 'id_of_the_algorithm::number_of_output', this syntax was used in line 9, algo_py_1::1, this represent that it will be passed as parameter the first output created by the algo_py_1, and the second parameter on line 10, is the number to be elevated to.

#### Supplementary Section 5: Bio-DIA user interface.

Access Bio-DIA through the link https://ucrania.imd.ufrn.br/biodia-app/, after that the user will see the login page (Figure S3A). In case the user does not have an account, he could create an account by clicking on the “subscribe” hyperlink and he will be sent to fill out the form (Figure S3B)

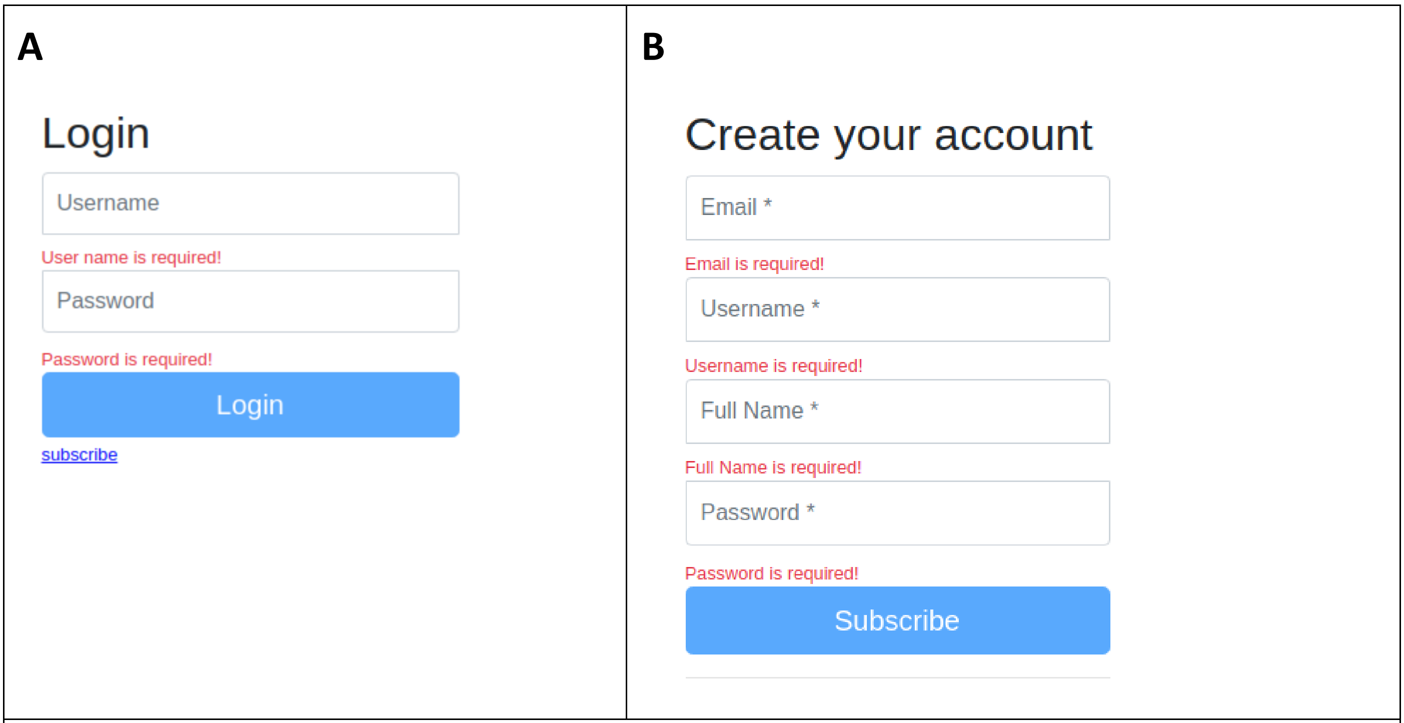


**Figure S3**: Log into Bio-DIA user interface.

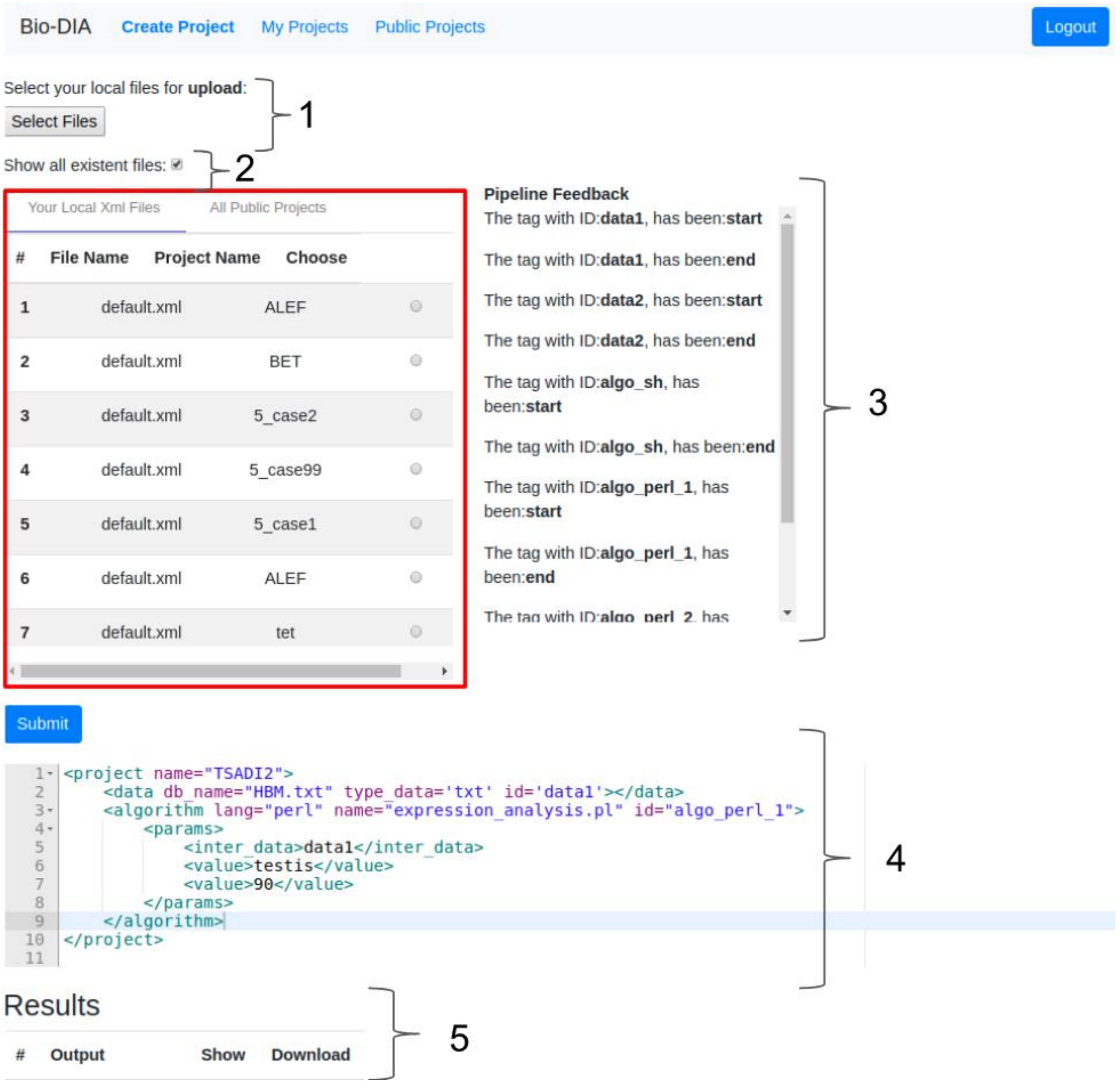


**Figure S4:** Bio-DIA main page. Creating a project. The braces group graphic elements of the interface to better explain them. Brace 1, to upload databases files such as: csv, tsv, or pipeline projects (XML files). After uploading your files, by checking the box pointed by the brace 2, the user can see and select all the XML files shown in the red rectangle. Brace 3 points to a frame in which are shown real time feedback messages from what is happening during the execution of every step of the project shown in brace 4. Brace 5 shows of results generated by the project.

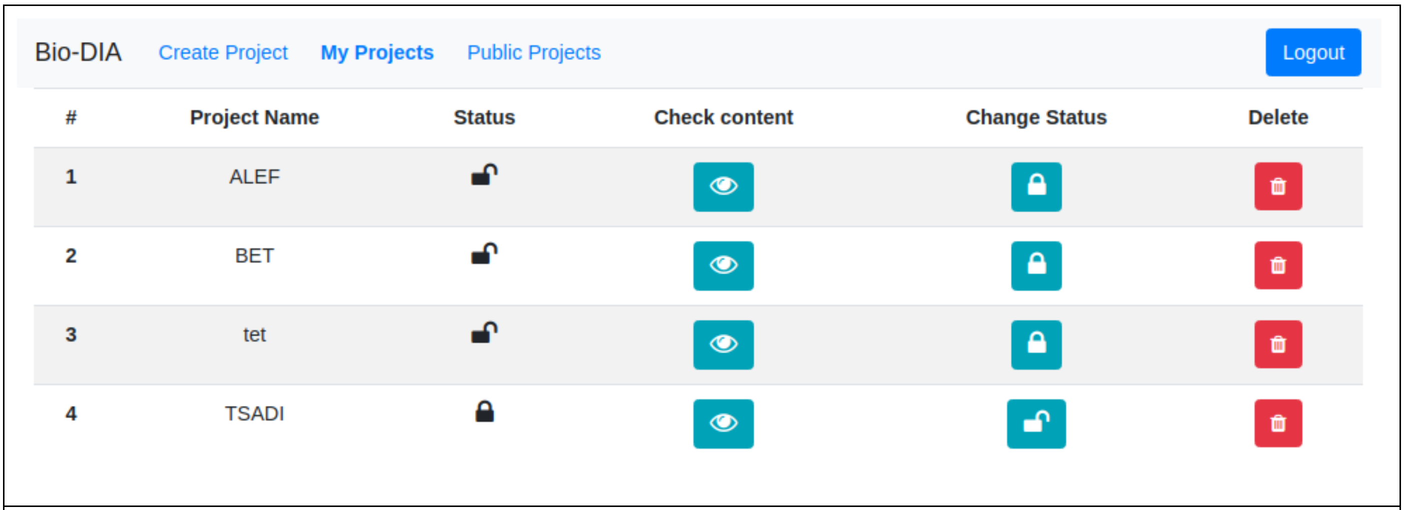


**Figure S5:** Bio-DIA project management page. This page allows the user to manage his projects. The first column "Project Name" list the projects names; the second column "Status" shows whether your project is public or private; the next column "Check content", clicking on the eye icon will show all the files that belongs to the project; the column "Change status" change the project status to private if the project is public, and vice versa; the last column 'Delete', remove the project and delete all the outputs created.

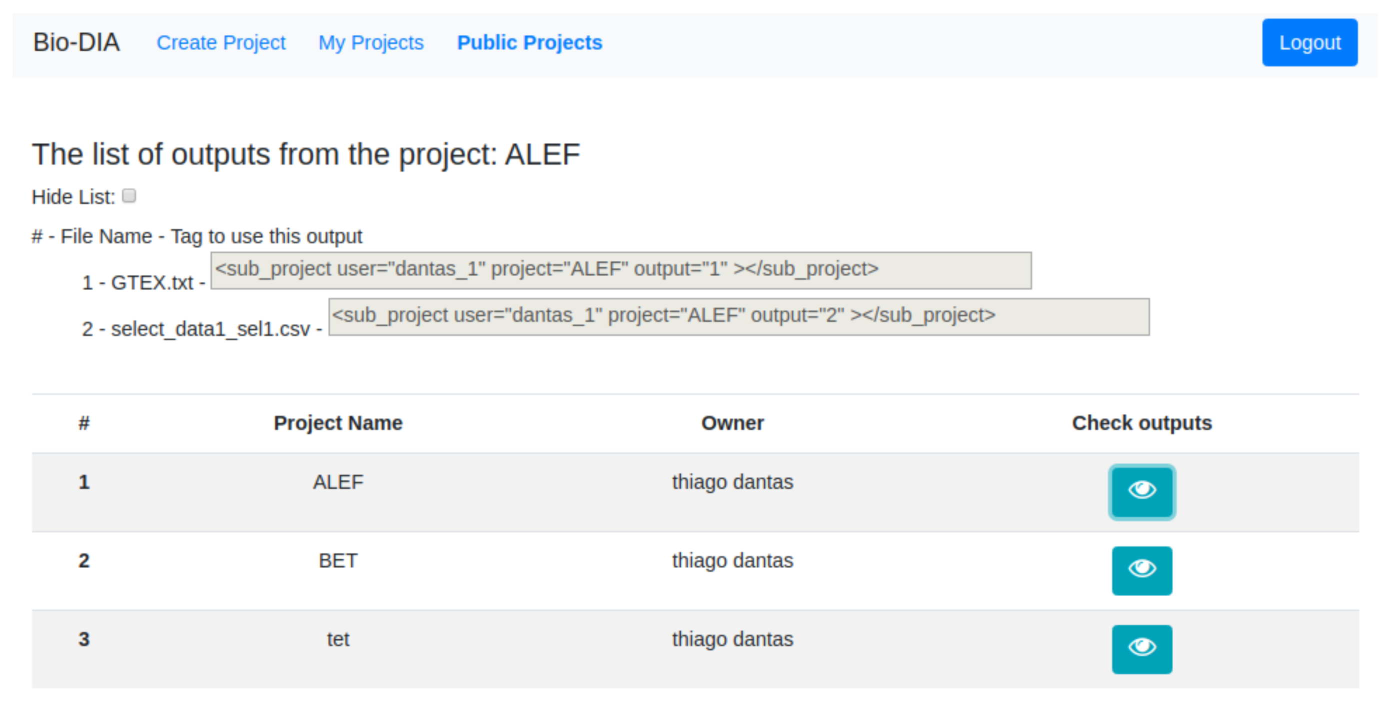


**Figure S6:** Bio-DIA public project page. This page lists all public projects of the system; hence all users can use and reproduce them. The project list shows the project name, its owner and by clicking on the eye icon at the "Check output", all outputs created by the project will appear above and the user just need to copy and paste the tag "sub_project" (gray text box) on their own pipeline to reuse the specific output.

#### Supplementary Section 6: Project examples

Log into Bio-DIA by access the link: https://ucrania.imd.ufrn.br/biodia-app/, make the login using, **login**: bioguest, **password**: welcome123. Copy this pipeline below and press the **submit** button.

**A**

```
1  <project name="pipe1">
2      <data db_name="GTEX.txt" type_data='txt' sep='\t' comment='#' id='data1'></data>
3      <select from="data1" id="sel1" get="vagina,lung,Gene Name" filter="lung == 19" limit='2'></select>
4      <save store="sel1" id="save2"></save>
5  </project>
```

**B**

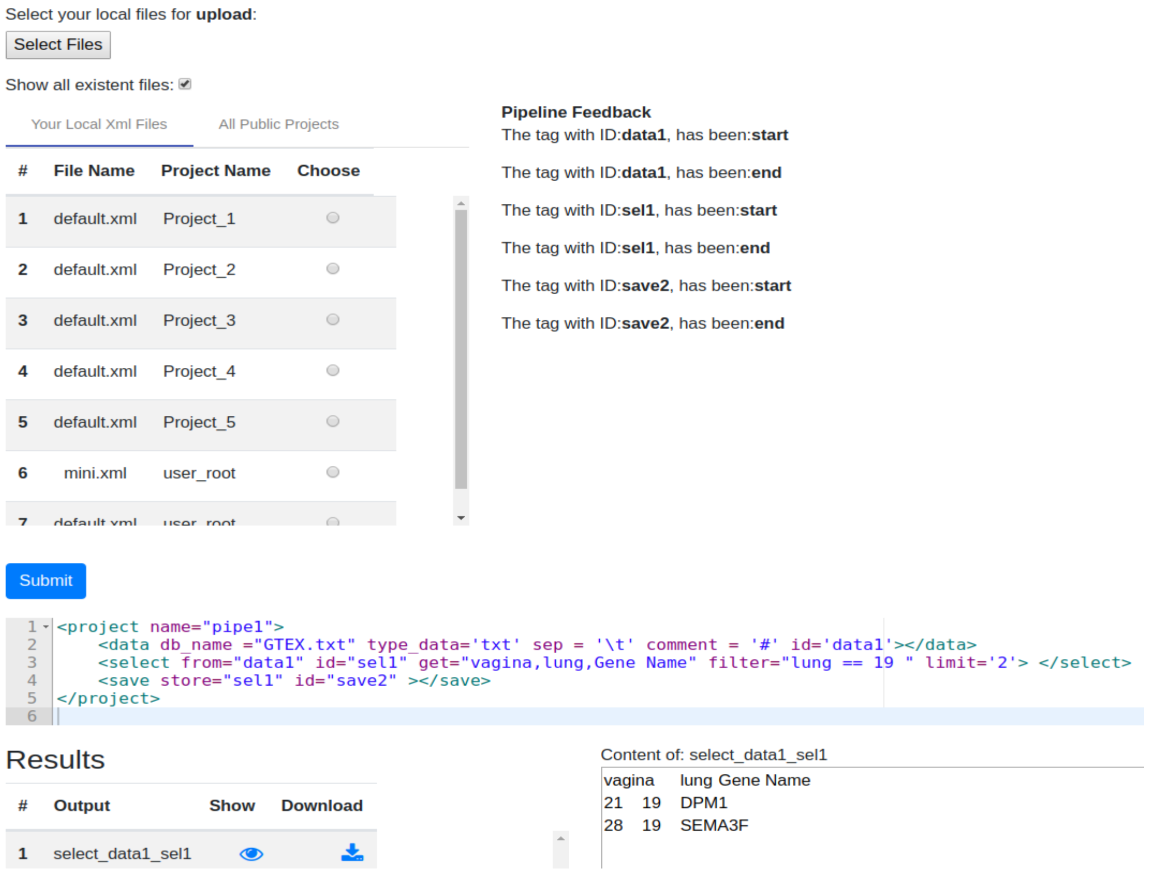


**Figure S7**: Simple pipeline to get started with Bio-DIA. This pipeline will load the file 'GTEX.txt' (line 2), which is a public file for all users. On the line 3 it will make a select on the file, getting 3 columns (vagina, lung and Gene Name), retrieving only the records where the lung values are equal to 19. In line 4, the result of this select is being saved on the project folder 'pipe1'. After finishing the pipeline on the Figure S7. Panel B shows the result page.

```
1   <project name="projectName">
2      <data db_name="sql_db" db_table="my_table" type_data="sql" id="data1"></data>
3      <select from="data1" select="myColumn, mySecondColumn" filter="myColumn > 2" id="select1"></select>
4      <algorithm lang="lang_of_script" name="script.extension" id="algorithm1">
5         <params>
6            <inter_data>data1</inter_data>
7            <value>my_static_value</value>
8         </params>
9      </algorithm>
10     <save store="select1" folder="myFolderName" id="save1"></save>
11  </project>
```

**Figure S8**: Project example using Bio-Dia XML tags. The parameter values have generic values to better explain the XML tags. The line 1 uses the tag project to create the project. For Bio-DIA, this is always the root tag. Line 2 is to connect with data; a table in a database in this case. Line 3, through a <select> tag, executes a selection and projection on data1 (it could be text files or tables in a relational databases). Line 4, through <algorithm> tag, executes scripts from languages (Shell, Python, Rscript or Perl). To pass the parameters to an algorithm an <params> tag embedded is used. Line 6 and 7 show two types of parameters: line 6 is using <inter_data> tag with a value making a reference to some id (data1), which was created previously in the project (Line 2) as an attribute of <data> or <select> tags; line 7 is using to pass static values. The line 10 is responsible, through <save> tag, to save in a specific folder data generated previously in the project. Line 10 is saving the content that makes reference to select1 (store="select1").

```
                              1                          2                  3
1   <sub_project user="userName_2" project="projectName" output="2" id="sub1" ></sub_project>
2   <save store="data1" folder="myFolderName" id="save1"></save>
```

**Figure S9:** How to reuse other project outputs. The <sub_project> tag allows to load results/outputs from others project previously executed. Line 1 is loading the project="projectName" (created in Fig. S1), that is owned by the user="userName_2", and the point 3 is the ID of the output that was generated by the project on point 2. For each output needed from other project, a <sub_project> tag must be created.
